# Predictive toxicology of chemical mixtures using proteome-wide thermal profiling and protein target properties

**DOI:** 10.1101/2023.07.04.547628

**Authors:** Veronica Lizano-Fallas, Ana Carrasco del Amor, Susana Cristobal

**Affiliations:** Department of Biomedical and Clinical Sciences, Cell Biology, Faculty of Medicine, Linköping University, Linköping 581 85, Sweden; Ikerbasque, Basque Foundation for Sciences, Department of Physiology, Faculty of Medicine, and Nursing, University of the Basque Country UPV/EHU, Leioa 489 40, Spain

**Author notes:** Corresponding Authors. Prof. Susana Cristobal. Department of Biomedical and Clinical Sciences, Cell Biology, Faculty of Medicine, Linköping University, Linköping, 581 85, Sweden.

**Keywords:** Proteome integral solubility alteration, thermal proteome profiling, chemical mixtures, exposome, predictive toxicology, TCDD, alpha-endosulfan, BPA

## Abstract

Our capability to predict the impact of exposure to chemical mixtures on environmental and human health is limited in comparison to the advances on the chemical characterization of the exposome. Current approaches, such as new approach methodologies, rely on the chemical mixture characterization and the available toxicological knowledge of individual compounds or similar mixtures. In this study, we show a new methodological approach for assessment of chemical mixtures based on a proteome-wide identification of the protein targets and revealing the relevance of new targets based on their role in the cellular crosstalk. We applied a proteome integral solubility alteration assay to identify 24 protein targets from a chemical mixture of 2,3,7,8- tetrachlorodibenzo-*p-*dioxin, alpha-endosulfan, and bisphenol A among the HepG2 soluble proteome, and validated the chemical mixture-target interaction orthogonally. To define the range of interactive capability of the new targets, the data from intrinsic properties of the targets were retrieved. Introducing the target properties as criteria for a multi-criteria decision-making analysis called the analytical hierarchy process, the prioritization of targets was based on their involvement in multiple pathways. This methodological approach that we present here opens a more realistic and achievable scenario to address the impact of complex and uncharacterized chemical mixtures in biological systems.

**SYNOPSIS:** Methods for unbiased identification of protein targets of chemical mixtures and prediction of their impact in human and environmental health.

## 1. Introduction

We are continuously exposed to chemical mixtures through sources such as air, water, food, and consumables. However, assessing the potential health effects of exposure to unknown mixtures poses significant challenges. To address this risk, the concept of “exposome” was introduced, initially focusing on evaluating the impact of environmental factors throughout an individual’s lifetime^1^. This concept has since expanded to include the measurement of effects on health resulting from cumulative exposures across a person’s life, encompassing external exposures, personal biology, and sociological factors^2^. Numerous studies in the field have concentrated on different aspects, such as investigating early-life exposures in population-based cohorts^3^, examining occupational exposures^4^, exploring the urban exposome^5^, or evaluating the impact on specific diseases^6^. Predicting adverse health outcomes from exposure to unknown chemical mixtures is still a significant challenge of the human exposome. Regulatory agencies and public health authorities recognize the lack of understanding regarding the potential effects of exposure to chemical mixtures, highlighting the pressing need for tools and methods to enhance our comprehension of the chemical mode of action within the context of mixture exposures^7^.

The new approach methodologies, such as *in vitro* methods, omics, quantitative structure-activity relationship, threshold of toxicological concern, read-across, toxicokinetic models, and *in silico* predictions, have been proposed as the emerging tools for assessing the hazard of mixtures^7–9^. Frameworks like the adverse outcome pathway (AOP) have also been developed to integrate knowledge from molecular, cellular, tissue, and organismal levels, aiding in the understanding of exposures to mixtures^10^. However, in a realistic scenario, our exposome at any given spatial and temporal point consists of hundreds of compounds in complex stoichiometry^11^. The chemical characterization of the studied mixtures that is still very difficult^12^, is a common requirement for the application of those new approach methodologies mixtures. Moreover, the requirement of preexisting toxicological knowledge of the individual compounds or similar mixtures are additional requirements add a first level of biasness to the applications. The second level of bias relates to the interpretation of the biological pathways that could be altered by mixtures based on expertise knowledge. As a result, studies focusing on unknown mixtures are scarce^13^, and the identification of underlying mechanisms for the effects of these mixtures remains unresolved^14^.

To address the first level of bias and overcome the challenge of characterizing mixtures, we recently proposed shifting the focus toward understanding the interaction between proteins and chemicals within the mixture. We suggested the application of proteomics-based methods for the unbiased identification of proteome-wide targets of chemical mixtures^15^. This approach draws parallels with the problem of drug-target engagement, where the specific targets of a novel drug need to be identified and characterized.

Several proteomics-based methods have already been developed to identify proteome-wide targets of drugs without bias. The concept originated from the use of thermal shift principles at a proteome scale, as described in the cellular thermal shift assay method^16^. Subsequently, the thermal proteome profiling method incorporated mass spectrometry to analyze the thermal shift assay, enabling high throughput and specificity in target identification^17^. Modifications were made to adapt this method for the analysis of novel bioactive compounds, which require a soluble proteome free from microsomal vesicles during the thermal shift assay^18^. Taking the analysis one step further, the proteome integral solubility alteration (PISA) method eliminates the need for sigmoidal fitting of protein melting curves, significantly increasing the throughput of the analysis by several orders of magnitude. The methodology has been developed to investigate potential drug targets from the entire soluble proteome and has been successfully used to validate known drug targets^19^. In our laboratory, we applied this method as a tool for assessing chemical mixtures with zebrafish embryo proteome as simplified organismal model^15^.

To address the second level of bias in assessing uncharacterized mixtures, we propose utilizing the two-dimensional PISA (2D PISA) method, which incorporates different concentrations of the chemical mixture, as it would enhance the robustness of identified targets^19^. Additionally, we combine the results from the 2D PISA method with the analytical hierarchy process (AHP) to determine which protein targets would have higher impact on cellular function disturbance. The AHP technique is a multi-criteria decision-making analysis tool used for ranking problems. It provides high-quality outputs and requires a moderate modeling effort, making it suitable for our application. In AHP, decision alternatives are ranked by considering key factors and assessing the relative value of each alternative target, incorporating evidence-based data^20, 21^. While expertise knowledge is valuable, this approach minimizes reliance on preexisting mechanistic understanding of individual chemicals within the mixture. Multi-criteria decision-making techniques have previously been used for various applications, such as prioritizing among different dust sources based on toxicity^22^, ranking chemicals for toxicological impact assessments^23, 24^, assessing the risk from multi-ingredient dietary supplements^25^, and integrating PISA data of individual chemicals into the AOP framework^26^. However, to the extent of our knowledge, this approach has not been previously applied to assessing unknown chemical mixtures. By combining the 2D PISA method with the AHP technique, we aim to overcome the limitations of previous approaches and provide a novel methodology for assessing the effects of unknown chemical mixtures.

The terminology surrounding exposure to chemical mixtures can often be imprecise. Therefore, it is necessary to clarify that in this study, we discuss chemical mixtures as the combined exposure to several chemicals, independent of their sources and routes of entry^27^. The concrete scope of this study is elucidating how chemical mixtures interact with proteins in a cellular native environment and using this information to predict the network of biological pathways that may be affected.

In this study, we apply the PISA method to identify a combination of protein targets from a chemical mixture. Additionally, we introduce the AHP method to determine which protein targets would have a higher impact on the disturbance of cellular function based on the intrinsic properties of the proteins. This methodology does not aim to offer or require the deconvolution of specific targets for each chemical in the mixture. However, the target list and their contribution to several cellular functions will assist on predicting which are the primary molecular and cellular pathways affected by the interaction with chemical mixtures and the potential consequences at the cellular level.

Furthermore, we anticipate that our results can guide the selection of bioassays at higher biological levels and provide knowledge that would contribute to the framework of the AOP concept. By understanding the protein targets and their implications in cellular function, we can gain insights into the broader biological effects of chemical mixtures and their potential impact on various biological systems.

## 2. Materials and methods

### 2.1. Sample preparation

Reagents and medium were purchased from Sigma-Aldrich (Sant Louis, MO, USA) unless otherwise noted. PBS was purchased from Trevigen (Gaithersburg, MD, USA). HepG2 cells were grown until 70 % confluence in EMEM medium supplemented with 7 % fetal bovine serum (ATCC, Manassas, VA, USA), 1675 mM L-glutamine, 85 U/mL penicillin, and 85 µg/ml streptomycin. Cells were harvested and centrifugated at 340 g for 4 min at 4 ℃. Three washes were made with 30 mL of cold PBS. Resuspension and centrifugation at 340 g for 4 min at 4 ℃. Washed pellets were snap-frozen in liquid nitrogen and stored at −80 ℃ until lysis.

### 2.2. Selection of tested mixture and concentrations

The chemical mixture analyzed in this study was composed of three persistent organic pollutants and endocrine disrupting compounds (EDCs) with different mechanisms of action which accumulate in the liver. The corresponding highest mixture concentration tested was 25 nM TCDD, 1 μM alpha-endosulfan, and 50 μM BPA.

The rationale for the selection of the test compounds was to include xenobiotics of high relevance for human and environmental health, which were previously studied individually or as mixtures. Therefore, we included compounds from previous studies for *in vitro* exposures to the human hepatic cell line HepaRG. This mixture originally contained TCDD, alpha-endosulfan, and we added BPA to gain complexity. TCDD and alpha-endosulfan can be found in the environment as a mixture because dioxins may contaminate chlorinated industrial products, as pesticides^28^. Moreover, TCDD and BPA are commonly found in food and have been studied as mixture through dietary supplementation equivalent to tolerable daily intake^29^. Concentration selection was based on the translation of external intakes into internal doses in hepatic cells and lower concentrations reporting effects^28, 30–32^.

### 2.3. 2D-PISA assay

HepG2 cells were resuspended in ice-cold PBS supplemented with protease inhibitors (Clontech, Mountain View, CA, USA) and lysed in ice bath by sonication in cycles of 10 s on /5 s off for 1 min at 6–10 μm amplitude at 25 % intensity from an exponential ultrasonic horn of 3 mm in a Soniprep 150 MSE (MSE Ltd., Lower Sydenham, London, UK). The insoluble parts were sedimented by centrifugation at 100,000 g for 60 min at 4 °C^18^. Protein concentration was determined by BCA assay^33^. The soluble proteome was used to perform the 2D PISA assay, as described in Gaetani et al. ^19^ with some modifications. Briefly, the soluble proteome and the studied chemical were incubated for 10 min at 25 °C. The incubation was performed with the following mixture at 10 different concentrations: TCDD + alpha-endosulfan + BPA. The studied concentrations were 100, 90, 80, 70, 60, 50, 40, 30, 20 and 0 %. The highest concentration (100 %) corresponds to 25 nM TCDD, 1 μM alpha-endosulfan, 50 μM BPA. The control samples (0%) were incubated in the presence of the vehicle, dimethyl sulfoxide (DMSO), utilized for the solubilization of the compounds. Ten specific temperatures were selected for the thermal assay: 37, 42, 46, 49, 51, 53, 55, 58, 62 and 67 °C. These temperatures were selected to ensure that at least 90 % of the studied proteins have their melting point within this range^17^. Aliquots containing 10 μg of protein (one for each of the temperatures in the entire range covered in the thermal shift assay) were independently heated at the corresponding temperature for 3 min, followed by 3 min at room temperature. For each concentration, aliquots of all temperature points were pooled and centrifuged at 100,000 g for 20 min at 4 °C, to remove the proteins that had an alteration in solubility after the thermal shift assay. Supernatants from intermediate concentrations were combined. The collection of soluble fractions in the supernatants from the three conditions (control – 0 %, intermediate concentrations and highest concentration – 100 %) were processed using a general bottom-up proteomics workflow and the purified peptides were analyzed by label-free nano liquid chromatography-tandem mass spectrometry analysis (nLC-MS/MS)^18^. Three biological replicates were performed for each experiment.

### 2.4. Filter aided sample preparation (FASP)

The samples were digested following the FASP method. First, the protein samples corresponding to the supernatants after centrifugation were prepared with SDT buffer (2 % SDS, 100 mM Tris-HCl, pH 7.6 and 100 mM DTT), according to Wiśniewski et al. ^34^. To perform FASP, the samples were diluted with 200 μl of 8 M urea in 0.1 M Tris/ HCl, pH 8.5 (UA) in 30 kDa microcon centrifugal filter units. The filter units were centrifuged at 14,000 g for 15 min at 20 °C. The concentrated samples were diluted with 200 μl of UA and centrifuged at 14,000 g for 15 min at 20 °C. After discharging the flow-through 100 μl of 0.05 M iodoacetamide was added to the filter units, mixed for 1 min at 600 rpm on a thermo-mixer, and incubated static for 20 min in dark. The solution was drained by spinning the filter units at 14,000 g for 10 min. The filter units were washed three times with 100 μl buffer UA and centrifuged at 14,000 g for 15 min. The filter units were washed three times with 100 μl of 50 mM ammonium bicarbonate. Endopeptidase trypsin solution in the ratio 1:100 was prepared with 50 mM ammonium bicarbonate, dispensed, and mixed at 600 rpm in the thermomixer for 1 min. These units were then incubated in a wet chamber at 37 °C for about 16 h to achieve effective trypsinization. After 16 h of incubation, the filter units were transferred into new collection tubes. To recover the digested peptides, the tubes were centrifuged at 14,000 g for 10 min. Peptide recovery was completed by rinsing the filters with 50 μl of 0.5 M NaCl and collected by centrifugation. The samples were acidified with 10 % formic acid (FA) to achieve pH between 3 and 2. The desalting process was performed by reverse phase chromatography in C18 top tips using acetonitrile (ACN; 60 % v/v) with FA (0.1 % v/v) for elution, and vacuum dried to be stored at −80 °C till further analysis.

### 2.5. Nano LC-MS/MS analysis

The desalted peptides were reconstituted with 0.1 % FA in ultra-pure milli-Q water and the concentration was measured using a Nanodrop (Thermo Fischer Scientific, Waltham, MA, USA). Peptides were analyzed in a QExactive quadrupole-orbitrap mass spectrometer (Thermo Fischer Scientific). Samples were separated using an EASY nLC 1200 system (Thermo Fischer Scientific) and tryptic peptides were injected into a pre-column (Acclaim PepMap 100 Å, 75 um × 2 cm) and peptide separation was performed using an EASY-Spray C18 reversed-phase nano LC column (PepMap RSLC C18, 2 um, 100 Å, 75 um × 25 cm). A linear gradient of 6 to 40 % buffer B (0.1 % FA in ACN) against buffer A (0.1 % FA in water) during 78 min and 100 % buffer B against buffer A till 100 min, was carried out with a constant flow rate of 300 nl/min. Full scan MS spectra were recorded in the positive mode electrospray ionization with an ion spray voltage power frequency (pf) of 1.9 kV (kV), a radio frequency lens voltage of 60 and a capillary temperature of 275 °C, at a resolution of 30,000 and top 15 intense ions were selected for MS/MS under an isolation width of 1.2 m/z units. The MS/MS scans with higher energy collision dissociation fragmentation at normalized collision energy of 27 % to fragment the ions in the collision induced dissociation mode.

### 2.6. Peptide and protein identification and quantification

Proteome Discoverer (v2.1, Thermo Fischer Scientific) was used for protein identification and quantification. The MS/MS spectra (. raw files) were searched by Sequest HT against the Human database from Uniprot (UP000005640; 95,959 entries). Cysteine carbamidomethylation was used as static modification and methionine oxidation as dynamic modification for both identification and quantification. A maximum of 2 tryptic cleavages were allowed, the precursor and fragment mass tolerance were 10 ppm and 0.02 Da, respectively. Peptides with a false discovery rate of less than 0.01 and validation based on q-value were used as identified. The minimum peptide length considered was 6 and the FDR was set to 0.1. Proteins were quantified using the average of top three peptide MS1-areas, yielding raw protein abundances. Common contaminants like human keratin and bovine trypsin were also included in the database during the searches for minimizing false identifications. The mass spectrometry proteomics data have been deposited to the ProteomeXchange Consortium *via* the PRIDE^35^ partner repository with the dataset identifier PXD039244.

### 2.7. Analysis of 2D PISA assay for target identification

The 2D PISA assay measures the protein abundance from 3 biological replicates of 3 conditions (control – 0 %, intermediate concentrations and highest concentration – 100 % of the tested compound). Protein abundances from control and the highest concentration represent, for each protein, the integral of the area under its melting curve within the used temperature interval. If S_m_ is the value for the control condition and S_m_’ is the corresponding value for the highest concentration condition, then the PISA analogue of the melting temperature shift (ΔT_m_) is

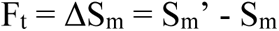

Protein abundance from intermediate concentrations (S_m_’’) represents an integral of the concentration-dependance curve. Similarly, the PISA analogue of the compound concentration required to induce half of the ΔT_m_ (C_0_) is

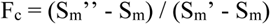

For each protein, the abundance was normalized on the average value for the control condition, and then F_t_ and F_c_ were calculated as described. Two-tailed Student’s t-test (with equal or unequal variance depending on F-test) was applied to calculate p-values. Proteins with p-values < 0.05 for both F_t_ and F_c_ were considered protein targets, meaning to be the proteins combining solubility alteration with action at a low compound concentration^19^. A cut-off value of 0.02 for both F_t_ and F_c_ p-values was applied to obtain the top protein targets. To facilitate the visualization of the protein targets, the data was represented in a scatter plot combining F_t_ and F_c_ p-values.

### 2.8. Nanoscale differential scanning fluorimetry (nanoDSF) analysis

One identified protein target was selected to perform the protein-mixture binding validation. We assessed the differential scanning fluorimetry with a nanoDSF device as an orthogonal validation approach. NanoDSF is based on the changes in the intrinsic tryptophan fluorescence resulting from alterations of the 3D-structure of proteins, when proteins unfold, as a function of the temperature. Therefore, a melting temperature can be determined^36^. Monitoring of the ITF at 330 nm and 350 nm during protein thermal denaturation was carried out in a Prometheus NT.48 instrument from NanoTemper Technologies (Munchen, Germany) with an excitation wavelength of 280 nm. Excitation power was set at 25 %. Capillaries were filled with 10 μl of a solution containing the protein and the mixture, placed into the sample holder and a temperature gradient of 0.5 °C/min from 20 °C to 80 °C was applied. The ratio of the recorded emission intensities (Em350nm/Em330nm), which represents the change in tryptophan fluorescence intensity was plotted as a function of the temperature. The fluorescence intensity ratio and its first derivative were calculated with the manufacturer’s software (PR.ThermControl, version 2.3.1 from NanoTemper Technologies). For validating protein-mixture binding, purified protein was mixed with 3 different concentrations of the mixture. The tested mixture concentrations were 20 %, 60 %, 70 %, 80 %, 90 % and 100 %, where 100 % corresponds 25 nM TCDD, 1 μM alpha-endosulfan, 50 μM BPA. Protein final concentration was 0.7 mg/ml. Control was performed with purified proteins in PBS and DMSO maintaining the corresponding protein concentration as for the mixture. Three replicates were carried out for each condition. Protein targets selection was based on the availability on the market (full-length recombinant protein and without GST tag, due to possible interferences with chemical binding) and the number of tryptophan residues (at least 2).

### 2.9. AHP analysis

The AHP approach arranges the factors considered to decide on a hierarchic structure and relies on three steps^21^. The first one is decomposition. Here, the problem was structured as a hierarchy, where the first level contains the overall goal, i.e., ranking the top targets according to disturbance of cellular function. The following level corresponds to the criteria which contribute to the goal in our case, criteria related to intrinsic properties of the protein targets. The third level of the hierarchy includes the alternatives (protein targets) to be evaluated in terms of the criteria in the second level. The second step is the elicitation of pairwise comparison judgments, where a matrix of the relative importance of each criterion over each other was performed using a scale from 1 to 9, according to the expertise of the authors. In this scale, 1 denotes that the two factors contribute equally to the goal, 9 represents the extreme importance of one over another, while 3 indicates slight importance. A numeric scale of 5 represents moderate importance and 7 indicates a very strong relevance of one factor over another. The values 2, 4, 6, and 8 represent the intermediate values between two adjacent judgments. After calculating the priority vector of the matrix, the consistency of the pairwise comparisons was evaluated through the calculation of the consistency ratio. A value of up to 10 % is considered acceptable^21^. The third step of AHP is to establish the global priorities of the alternatives. This was done by laying out the local priorities of the alternatives concerning each criterion in a matrix (by pairwise comparison judgments using a scale from 1 to 9). Equations for the calculations are described in detail in our previous study^26^. The alternative with the highest global priority was ranked first, meaning that proteins with lower number in the rank could produce a higher disturbance of cellular function when they bind to the mixture. After the ranking of the targets was obtained through the AHP technique, a sensitivity analysis was performed to assess the stability of the optimal solution under variations in the parameters. Therefore, the most critical criterion and the smallest change in the current weight of a criterion that modifies the ranking of the alternatives are determined^37^. The priority vector, global priorities (ranking) of the alternatives, consistency ratios, and sensitive analysis were calculated using the online software AHP-OS^38^.

### 2.10. Analysis of biological pathways and their networks

The top targets identified by the 2D PISA assay were subjected to a pathway over-representation analysis in Reactome v82^39^. Enriched pathways with a p-value equal or lower than 0.05 involving targets with a lower position in the AHP rank were considered for the prediction of the network of biological pathways that could be altered by the protein-mixture interaction.

## 3. Results

### 3.1. Identification of protein targets from a chemical mixture in hepatic cells

To identify the proteins that interact with a chemical mixture composed of TCDD, alpha-endosulfan, and BPA in the soluble proteome from hepatocytes we applied the 2D PISA assay. This proteome-based thermal shift assay detects chemical-induced solubility changes in the protein targets. The assay was conducted using a temperature-based approach at 10 different concentrations of the chemical mixture. After analyzing the soluble proteome, we identified and quantified a total of 2,886 proteins across the biological replicates. Furthermore, 1,493 proteins were quantified in all three replicates and subsequently subjected to the 2D PISA assay. Among these, 24 proteins exhibited significant alterations in solubility in both temperature (F_t_) and concentration (F_c_) dimensions, thus identifying them as targets (Supplementary table 1).

The top protein targets were glycine-tRNA ligase (GARS), leukotriene A-4 hydrolase (LTA4H), ubiquitin carboxyl-terminal hydrolase (UCHL5), RNA-binding protein FUS (FUS), RUN and FYVE domain-containing protein 1 (RUFY1), alpha-actinin-4 (ACTN4), protein CDV3 homolog (CDV3), lupus La protein (SSB), peroxiredoxin-5 (PRDX5) and lysosomal alpha-glucosidase (GAA) (Figure 1).

**Figure 1.**
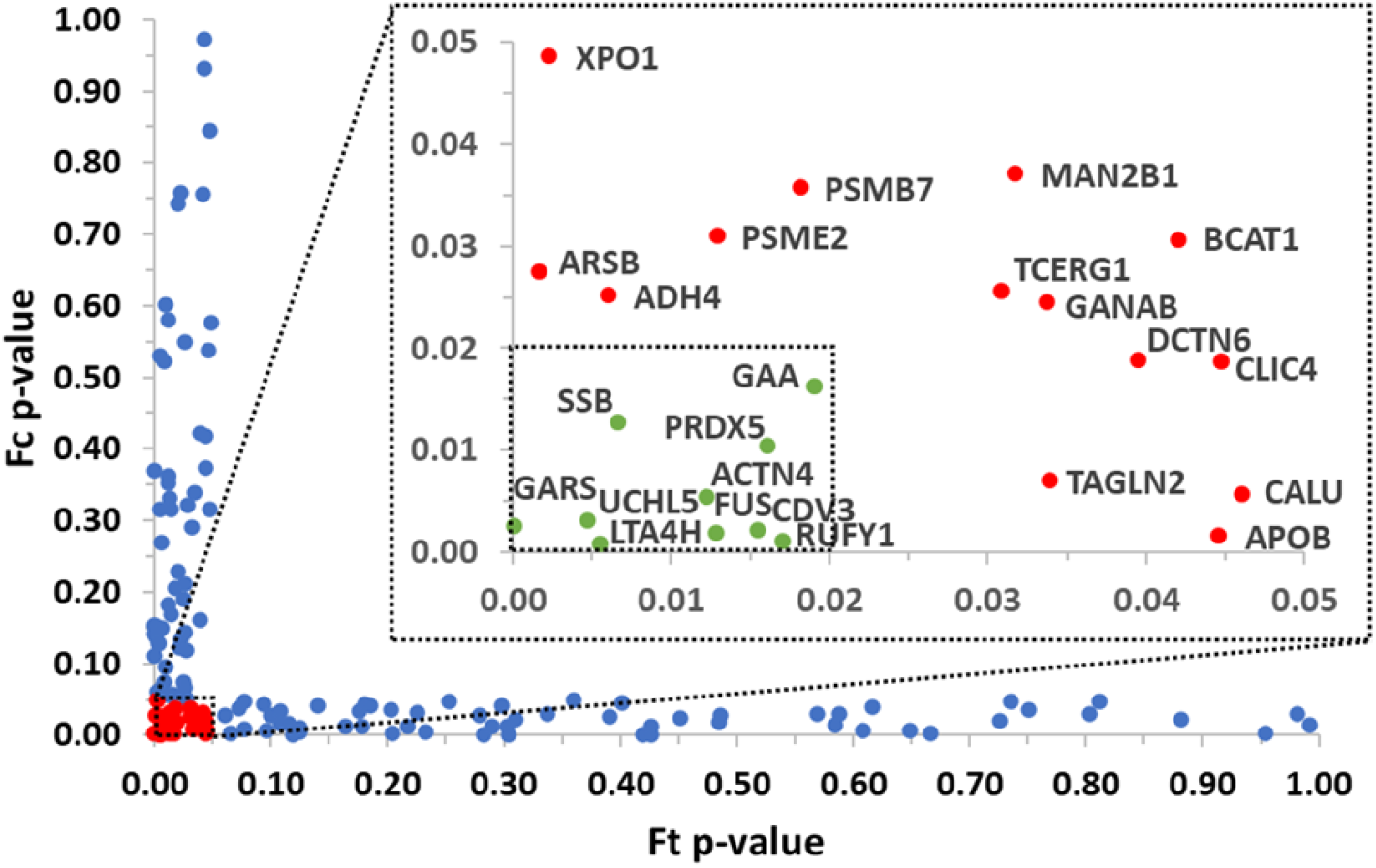
Protein targets of the studied mixture in the soluble proteome from hepatocytes identified by 2D PISA assay. The studied concentrations were 100, 90, 80, 70, 60, 50, 40, 30, 20, and 0 %. The highest concentration (100 %) corresponds to 25 nM TCDD, 1 uM alpha-endosulfan, and 50 uM BPA. The significant proteins for F_t_ or F_c_ are shown in blue. The identified protein targets are shown in red and labeled and the top protein targets are shown in green. Ft: PISA analogue of the melting temperature shift, and Fc: PISA analogue of the compound concentration required to induce half of the melting temperature shift.

### 3.2. Structural validation of the interaction between chemicals and targets

Next, our aim was to validate the direct interaction between the targets identified through PISA and the chemical mixture. To achieve this, we selected a purified target protein and performed nanoDSF a broad range of chemical concentrations of the mixture to evaluate the effects of the interaction at the structural level. This method serves as a suitable approach for validating protein-mixture binding, as it enables the detection of modifications in protein melting temperature resulting from changes in protein stability due to interactions with the chemical mixture.

To conduct the binding validation using nanoDSF, we chose the protein target chloride intracellular channel protein 4 (CLIC4) from the results of the 2D PISA assay. CLIC4 was an ideal candidate for this analysis as it possesses 2 tryptophan residues, a requirement for the nanoDSF technique. The full-length recombinant protein with an N-Terminal His-tag (NBP1-51032) was procured from Novus Biologicals (Centennial, CO, USA) for this purpose.

Following the nanoDSF validation, we conducted a comparison between different concentrations (20 %, 60 %, and 100 %) of the chemical mixture against the control. Among these, only the interaction between CLIC4 and the mixture at a concentration of 100 % exhibited a noticeable shift in the melting temperature (averaging from 55.4 °C to 47.2 °C) as depicted in Figure 2A.

**Figure 2.**
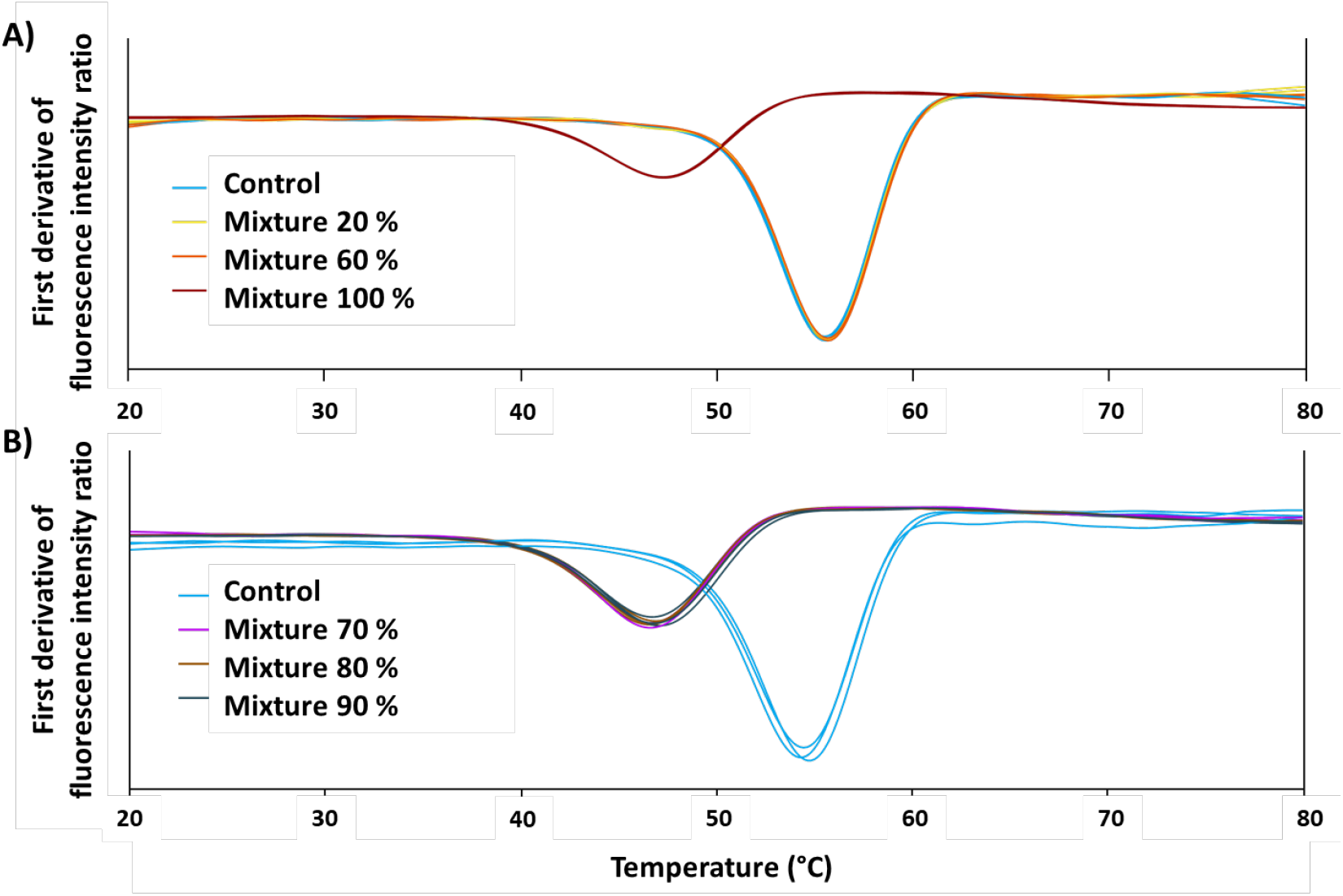
NanoDSF analysis of protein target CLIC4 exposed to the studied mixture of chemicals. The graphs showed the alteration in melting temperatures from CLIC4 with different concentrations of the mixture: **A)** 20 % (yellow), 60 % (orange), and 100 % (red), and **B)** 70 % (purple), 80 % (brown), and 90 % (dark blue), where a 100 % concentration corresponds to 25 nM TCDD, 1 uM alpha-endosulfan, and 50 uM BPA. A control without the mixture was included in both experiments (light blue).

To further verify the binding between the selected target and the mixture at intermediate concentrations, a second experiment was performed utilizing concentrations of 70 %, 80 %, and 90 %. In comparison to the control, a consistent shift in the melting temperature (averaging from 54.4 °C to 46.7 °C) was observed across all three intermediate concentrations tested, as shown in Figure 2B.

### 3.3. Evaluating the significance of identified protein targets in cellular crosstalk

We have applied the AHP method to rank the top targets, identified by the 2D PISA assay, according to the disturbance of cellular function their impairment could cause. The first challenge for the application of AHP was to generate a hierarchical structure of the problem. The first level of the hierarchy was the overall goal of the analysis, i.e., ranking the top targets according to disturbance of cellular function. The second level contained 7 criteria that were selected based on the need to predict the impact of chemical mixtures on human health when the characterization of the mixture is lacking. Those criteria are based on the intrinsic properties of the proteins that will determine the cellular function, and cellular crosstalk, ranging from structure to functional specificity in tissues. Criteria are described in Table 1. The third level included the top protein targets identified by the 2D PISA assay as the alternatives to be evaluated in terms of the criteria of the second level.

**Table 1.** Description of the selected criteria which contribute to the overall goal of ranking by AHP the top targets identified for the chemical mixture by 2D PISA assay.

| Criteria | Source | Description | Selection support |
| --- | --- | --- | --- |
| **Position in $F_t$ (solubility alteration) ranking** | 2D PISA assay | Protein with the highest solubility alteration (the lowest position in the ranking) is more likely to bind the mixture. Disturbance is not expected if there is no binding. | (Gaetani et al., 2019; Lizano-Fallas et al., 2021) |
| **Position in $F_c$ p-value ranking** | 2D PISA assay | Protein with the highest significance (the lowest position in the ranking) for solubility alteration is more likely to bind the mixture. Disturbance is not expected if there is no binding. | (Gaetani et al., 2019; Lizano-Fallas et al., 2021) |
| **Protein subcellular localization** | UniProt (Release 2022_05) (The UniProt Consortium, 2023)<br>UniProt annotation was used. Repetitive locations were not included. | Protein with the highest number of subcellular localizations should have an important function and compromise more organelles functions, therefore a higher disturbance is expected. | (Bauer et al., 2015; Thul et al., 2017) |
| **Protein structural domains** | UniProt (Release 2022_05)<br>Pfam entries were used. Only the entries with domain as Pfam type were included. Repeated domains were included. | Protein with the highest number of structural domains relates with more connections and biological functionalities, therefore a higher disturbance is expected. | (Chothia et al., 2003; Middleton et al., 2010; Wang et al., 2021) |
| **Protein abundance** | PAXdb (4.2)<br>(Wang et al., 2015)<br>Abundance in ppm for the whole organism (*Homo sapiens*) was used. | Protein with the highest abundance tends to have more protein-protein interactions and correlate with essentiality, therefore a higher disturbance is expected. | (Berggård et al., 2007; Ishihama et al., 2008) |
| **Protein hydrophobic surface** | Hi-patch (v1)<br>(Sánchez-Morán et al., 2021) | Protein with the highest number of hydrophobic surface patches relates to protein-protein interactions, membrane binding, and progression of diseases, therefore a higher disturbance is expected. | (Malleshappa Gowder et al., 2014; van Gils et al., 2022; Young et al., 1994) |
| **Tissue specificity** | Human Protein Atlas (22.0) | Protein with preferential functions in the studied tissue is expected to show a higher disturbance. | (Jiang et al., 2020; Uhlén et al., 2015) |

After defining the 7 criteria for ranking the top targets, a matrix of pairwise comparison judgments of the criteria was performed by the authors’ expertise. Table 2 shows the obtained matrix, the priority vector, and the corresponding consistency ratio. According to the priority vector, the criteria: position in F_t_ (solubility alteration) ranking and position in F_c_ p-value ranking obtained the highest weight of relevance.

**Table 2.**
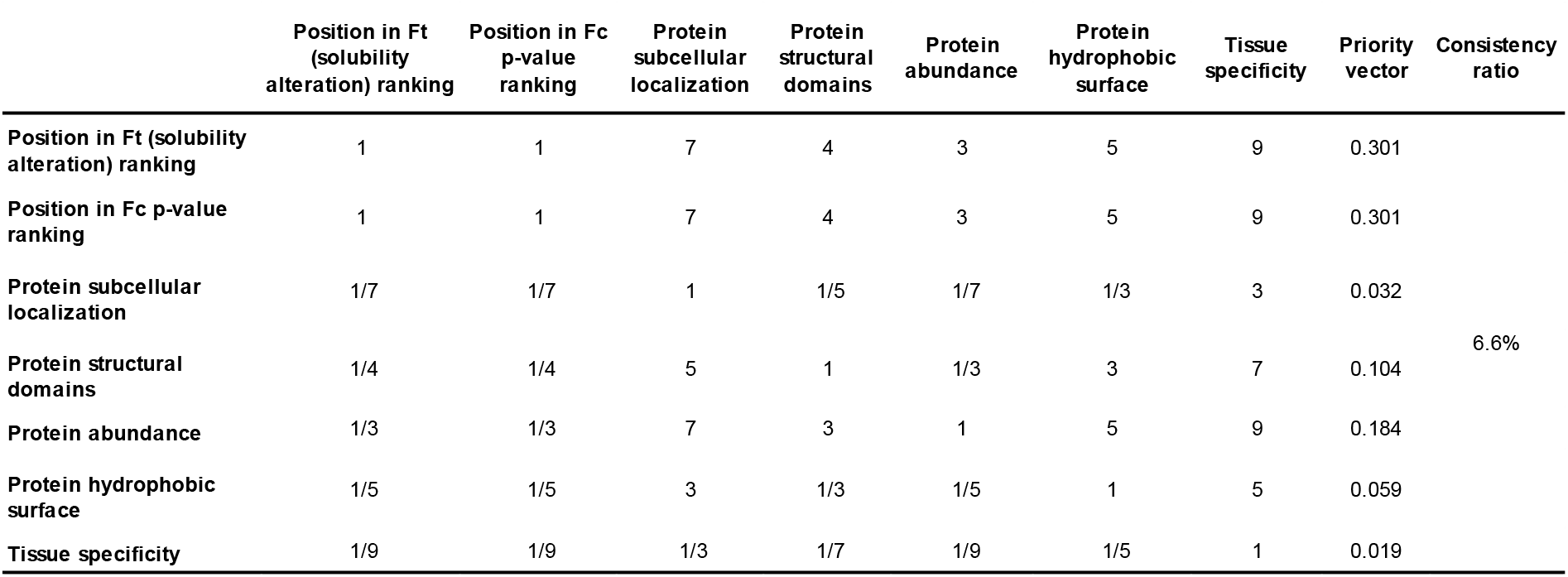
Pairwise comparison matrix for the criteria at level 2 of the hierarchy and the computed values of priority vector and consistency ratio.

The following criteria in the priority rank were: i) protein abundance, ii) protein structural domains, and iii) protein hydrophobic surface. Finally, the lowest weight was occupied by the criterion tissue specificity.

The third step of AHP involves determining the global priorities of the alternatives through pairwise comparison judgments. To facilitate this process, a dedicated database was created, containing information on each protein for each criterion (Supplementary table 2). The sources used to compile this information are outlined in Table 1. The matrices of pairwise comparison judgments of the alternatives were performed, and the corresponding local priority vectors and consistency ratios are shown in Supplementary table 3. Global priorities were then established based on these assessments. The top-ranked targets were those with the highest global priority, reflecting their potential to cause significant disturbance to cellular function.

In the ranking obtained, the proteins CDV3, ACTN4, and LTA4H secured the first three positions, indicating their strong potential for cellular disturbance. On the other hand, the protein GAA ranked last in the hierarchy (Table 3).

**Table 3.** Global priority and ranking of the top protein targets (alternatives).

| <u>Protein</u> | <u>Global priority</u> | <u>Rank</u> |
| --- | --- | --- |
| CDV3 | 0.148 | 1 |
| ACTN4 | 0.133 | 2 |
| LTA4H | 0.127 | 3 |
| UCHL5 | 0.118 | 4 |
| FUS | 0.103 | 5 |
| RUFY1 | 0.094 | 6 |
| PRDX5 | 0.089 | 7 |
| SSB | 0.074 | 8 |
| GARS | 0.065 | 9 |
| GAA | 0.049 | 10 |

In addition, a sensitivity analysis was conducted to assess the robustness of the targets’ ranking derived from the AHP methodology. The criterion “position in F_c_ p-value” was identified as the critical factor, wherein even a light alteration in its weight can potentially impact the ranking order. Specifically, a relative change of 10.7 % in the weight assigned to this criterion can lead to a shift in the ranking between the proteins CDV3 and LTA4H, while a relative change of 2.3 % can impact the ranking between the alternatives ACTN4 and LTA4H. These findings highlight the sensitivity of the ranking outcome to variations in the weight assigned to the “position in F_c_ p- value” criterion.

### 3.4. Validating the prediction capability of the AHP ranking based on intrinsic properties of identified protein targets

The pathway enrichment analysis of the top targets showed 61 pathways under the threshold of a p-value of 0.05. The targets ACTN4 and LTA4H, which occupied the lower positions in the AHP analysis, are involved in 10 of these pathways (Figure 3). The protein CDV3, which has the first position in the ranking was not found in Reactome, however it has been considered for the prediction of the main molecular and cellular pathways that could be altered by the protein-mixture interaction. Therefore, in our study case, the prediction of the interference of the chemical mixture can be depicted as an alteration of biological pathways as cell proliferation; cell-cell communication through the nephrin family interactions; and metabolism of lipids through the biosynthesis of specialized proresolving mediators and synthesis of leukotrienes and eoxins.

**Figure 3.**
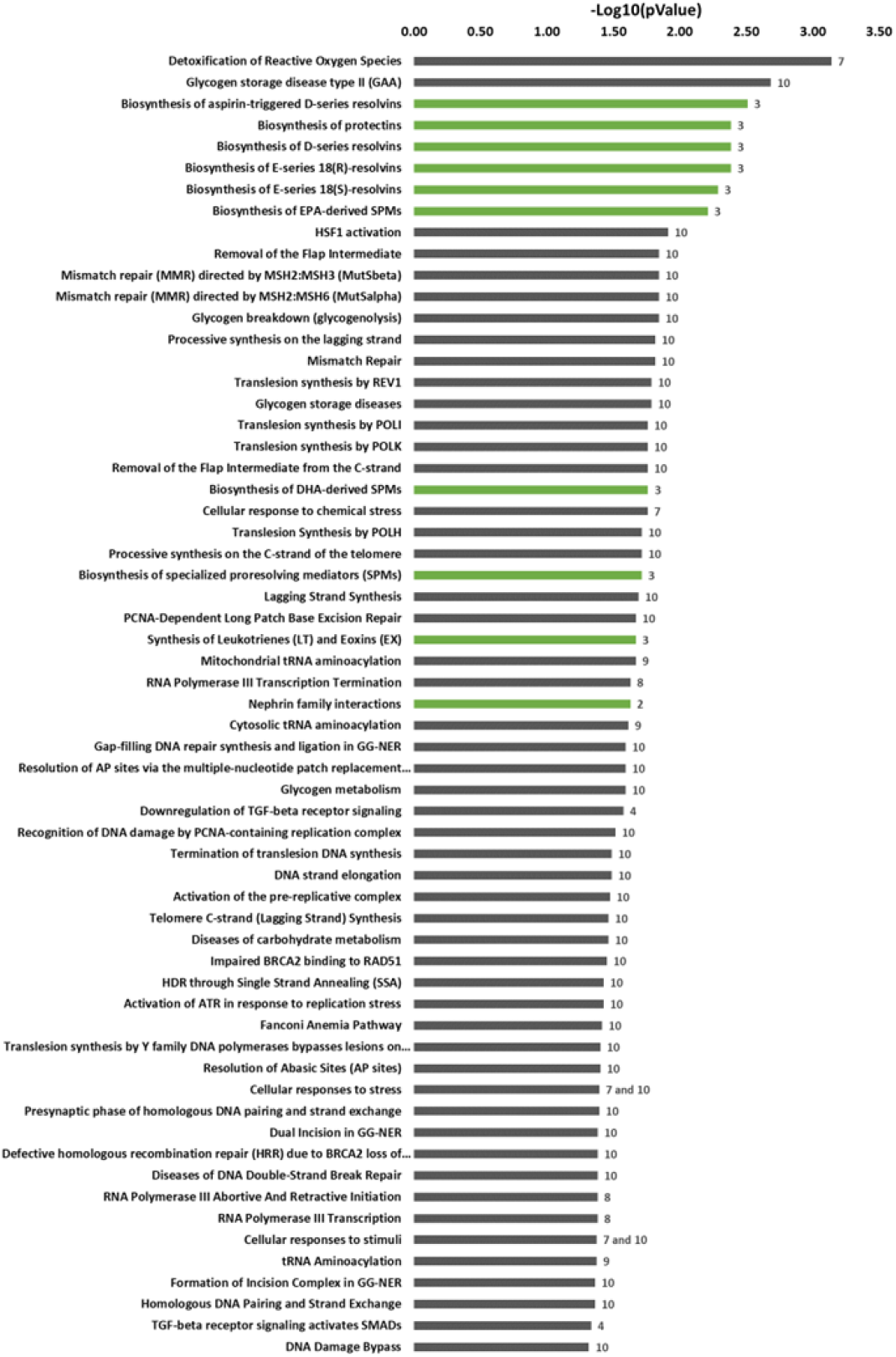
Pathway analysis showing significant enriched pathways for the top protein targets. The position in the AHP rank of the targets involved in each pathway is shown to the right side of each bar. The main molecular and cellular pathways that could be altered by the protein-mixture interaction are shown in green.

## 4. Discussion

Most of the complex mixtures of chemicals we encounter daily remain largely unstudied. To address this knowledge gap, it is crucial to focus on unbiased and high-throughput methodologies that do not only rely on detailed characterization or availability from toxicological data. In our study, we introduced a novel approach that combines the 2D PISA assay and the AHP technique. This method allows us to identify new protein targets within complex mixtures and rank them based on their intrinsic properties associated with cellular functionality. By focusing on the intrinsic properties of proteins, we gain a practical and feasible way to assess the impact of complex chemical mixtures on biological systems. This approach enhances our understanding of potential functional disturbances and provides comprehensive insights into the effects of these mixtures in a realistic manner.

The rationale behind thermal shift assay methods for identifying the target from ligands is based on how ligands could alter protein solubility and protein conformation. When a ligand or small molecule binds to a protein, it can induce conformational changes that may disrupt the normal folding pattern of the protein, leading to misfolding and subsequent aggregation^40^. Moreover, the conformational ensembles of protein monomers can exist in different states, and ligand binding often influences the conformation towards a specific set of ensembles to facilitate a particular function. Similarly, when considering protein oligomerization, the number of conformational ensembles must also be restricted. Therefore, the interaction of small molecules with a protein or a subunit of a protein oligomer can further modulate their conformational states and affect the overall structure and function. This interplay between ligand binding, conformational ensembles is crucial for understanding the intricate mechanisms underlying protein function and dysfunction^41^.

In previous studies, we successfully demonstrated the effectiveness of the PISA assay in identifying protein interactions from individual chemicals from cell lines^26^ and both single chemical and chemical mixture in zebrafish embryo model organism^15^. In this study, we opted for the 2D PISA approach, as opposed to the one-dimensional PISA utilized previously, to enhance the robustness of target identification. The 2D PISA assay offers an internal validation mechanism for the identified targets by considering significant changes in protein solubility across both dimensions (temperature and concentration). Using this method, a protein is designated as a target only if there is a significant alteration in its solubility in both dimensions. This selection criterion enhances the reliability and confidence of our target identification process.

Furthermore, we conducted an orthogonal validation of the binding between the protein mixture and our target proteins at a structural level by nanoDSF. Our focus was on confirming the existence of interactions between the protein and the mixture, rather than exploring the specific effects of these interactions. This method provides the changes in the melting temperature of proteins that is an indication of alterations in protein stability resulting from interactions with chemicals. While the 2D PISA assay involved pooling eight intermediate concentrations, it does not provide information about specific targets at each concentration. It is also important to note that the identification of a target at one intermediate concentration does not guarantee binding at all other tested concentrations. Nonetheless, we observed a shift in the melting temperature of the target protein CLIC4 at higher intermediate concentrations of the mixture. This observation serves as additional evidence confirming the structural binding between the protein mixture and our target protein. For this analysis, we studied purified protein targets, effectively eliminating the influence of cellular crowding effects that could potentially impact proteome-based analyses. Macromolecular crowding can indiscriminately enhance interactions and affect interaction rates^42^. However, it can also reduce the diffusion rate^43^. Eliminating these confounding factors, we could still confirm the protein-mixture interactions.

Our second aim was to introduce a method that could offer an unbiased selection or prioritization of the most relevant identified targets considering: the functional relevance to toxicology of mapping protein participating in networks, and availability to retrieve data for each individual protein target. The criteria based on the intrinsic properties of the proteins fulfilled those premises. The data for the selected criteria were either available in public databases or easily calculated without requiring deep knowledge in bioinformatics or programming. The data required for the selected criteria is readily available in public databases such as UniProt (The UniProt Consortium, 2023), PAXdb^44^, and The Human Protein Atlas ^45^, or can be easily calculated without the need for extensive bioinformatics or programming expertise ^46^. This accessibility facilitates the application of our approach for hazard characterization.

The main criteria tried to identify proteins capable of interacting with many other proteins and biomolecules in the cell. This type of protein, such as the hub proteins, due to their extensive interactions, could influence multiple cellular functions and are often involved in essential processes such as cell signaling, metabolism, and gene regulation. Previous studies have demonstrated that defects in hub proteins can lead to failures in a significant portion of the network, while defects in non-hub proteins have a less drastic impact on network function^47^. Moreover, hub proteins are known to play critical roles in the reproduction and survival of organisms compared to non-hub proteins^48^. For predictive toxicology, the possible disfunction of protein targets with larger number of connections would likely cause higher functional impact and therefore the focus for target prioritization, whereas in drug design the focus is to manipulate individual pathways causing minimal side-effects^49^.

After selecting the criteria for ranking the targets, a pairwise comparison was conducted to determine the weight or priority of each criterion. Among the seven chosen criteria, the positions in the F_t_ (solubility alteration) and F_c_ p-value rankings received the highest weight, indicating their significance in assessing the level of functional disturbance. Notably, if there is no binding between a chemical mixture and a protein, no disturbance is expected. The remaining five criteria were ordered in the priority vector based on their relevance from general to specific functional disturbance. The criterion of general protein abundance throughout the body carried the highest weight, while the criterion of specific preferential functions in the studied tissue (e.g., liver in this study) had the lowest weight. Later, the global priorities of the alternatives were established after performing pairwise comparison for each criterion. CDV3, ACTN4, and LTA4H ranked first, indicating a higher expectation of cellular disturbance resulting from their impairment. However, if the ranking had relied on the number of interactors for each protein, as derived from the IntAct database^50^, as seen in systems toxicology approaches^51^, the positions in the rank for these targets would be ninth, seventh, and third, respectively. This result underscores the relevance of utilizing an unbiased, comprehensive, and systematic approach that considers intrinsic protein properties as proxies for protein-protein interactions in complex networks. This approach avoids exclusion of targets that have not been extensively studied before, unlike methods solely based on preexisting knowledge of protein interactions.

Moreover, we anticipate that constructing the ranking of protein targets based on universally available intrinsic properties associated with cellular functionality would facilitate the extrapolation of toxicity predictions across species, even for non-model organisms. In this scenario, it would be feasible to predict the impact of chemical mixtures on any species solely using their genome/proteome database.

To validate our AHP results, a sensitivity analysis was conducted to assess the stability of the ranking obtained through the analysis. CDV3, ACTN4, and LTA4H, the top three targets, were closely positioned in terms of global priority values. Hence, their ranking positions may exhibit some variability. Consequently, these proteins were considered for predicting the key molecular and cellular pathways that could be affected by the protein-mixture interaction.

As an additional supporting strategy, we compared our predicted results with the reported effects of similar mixtures or their individual components. The chemical mixture used in this study consisted of 25 nM TCDD, 1 μM alpha-endosulfan, and 50 μM BPA. Literature findings supported our predictions, as effects on cell proliferation have been reported for BPA and endosulfan as individual compounds^52–54^. Similar effects have also been observed for mixtures of endocrine-disrupting compounds (EDCs) ^55^ and a hepatocyte cell line exposed to a mixture of TCDD and alpha-endosulfan, with comparable concentrations to those used in our study^30^. Regarding cell-cell communication, impairment in podocytes due to BPA exposure has been reported^56^. TCDD, endosulfan, and EDCs, in general, have been associated with effects in various tissues^57–60^. Furthermore, mixtures of persistent organic pollutants, including TCDD, and individual compounds such as endosulfan, have been linked to disruptions in lipid metabolism^55, 61^. While interactions between contaminants and established target drug families are well-known, the attention is increasingly shifting towards nonclassical targets like the ones presented in our study^62^, and understanding their characteristics has become important for making predictions and for prioritization.

## Conclusions

Our findings demonstrate that the 2D PISA assay effectively identifies protein targets from uncharacterized and complex mixtures with robustness. Furthermore, incorporating universally available intrinsic properties from the target proteins using the AHP technique enhances the accuracy of toxicology prediction, preventing the exclusion of unstudied targets. We envision that integrating high-throughput identification of mixture targets through proteomics-based thermal shift methods, along with target ranking using a semiquantitative AHP approach, has the potential to facilitate interspecies extrapolation of toxicity predictions, even for non-model organisms. This comprehensive approach holds promise for advancing the field of toxicology and improving our ability to assess mixture toxicity.

## Supporting information

Supplementary table 1. Protein targets identified after applying 2D PISA experiment to HepG2 cells soluble proteome exposed to the mixture

Supplementary table 2. Database containing the information of each protein target (alternatives) for each criterion.

Supplementary table 3. Pairwise comparison matrices for the alternatives at level 3 of the hierarchy for each criterion

## ABBREVIATIONS

AOP: adverse outcome pathway
PISA: the proteome integral solubility alteration
2D PISA: two-dimensional PISA
AHP: analytical hierarchy process
EDCs: endocrine disrupting compounds
DMSO: dimethyl sulfoxide
nLC-MS/MS: nano liquid chromatography-tandem mass spectrometry analysis
FASP: filter aided sample preparation
ΔT_m_: melting temperature shift
nanoDSF: nanoscale differential scanning fluorimetry
GARS: glycine-tRNA ligase
LTA4H: leukotriene A-4 hydrolase
UCHL5: ubiquitin carboxyl-terminal hydrolase
FUS: RNA-binding protein FUS
RUFY1: RUN and FYVE domain-containing protein 1
ACTN4: alpha-actinin-4
CDV3: protein CDV3 homolog
SSB: lupus La protein
PRDX5: peroxiredoxin-5
GAA: lysosomal alpha-glucosidase

## ASSOCIATED CONTENT

**Supplementary table 1**. Protein targets identified after applying 2D PISA experiment to HepG2 cells soluble proteome exposed to the mixture 25 nM TCDD 1 uM alpha-endosulfan 50 uM bisphenol A.

**Supplementary table 2.** Database containing the information of each protein target (alternatives) for each criterion.

**Supplementary table 3.** Pairwise comparison matrices for the alternatives at level 3 of the hierarchy for each criterion and the computed values of local priority vector and consistency ratio (CR) for each matrix.

## AUTHOR INFORMATION

### Author Contributions

VLF has performed most of the laboratory work and analyzed the data, contributed part of the analytical design, and wrote the first draft of the manuscript. ACA conducted some additional experiments. SC generated the idea and designed the study, supervised the analysis of the experimental work, discussed and interpretation of the results, wrote and modify the manuscript, edited the manuscript, and was responsible for funding acquisition. All the authors have contributed to discussions and modifications of the manuscript and approved it.

## Acknowledgments

All the mass spectrometry analysis has been performed with instrumentation from LiU MS facility. We thank Olatz Fresnedo from the University of the Basque Country UPV/EHU, Spain for kindly providing, the HepG2 cells.

## Data Availability Statement

The mass spectrometry proteomics data have been deposited to the ProteomeXchange Consortium via the PRIDE^35^ partner repository with the dataset identifier PXD039244.

